# Single molecule mechanics reveal Kif15 as an active molecular ratchet with acute strain sensitivity

**DOI:** 10.1101/141978

**Authors:** T. McHugh, H. Drechsler, A. D. McAinsh, N. J. Carter, R. A. Cross

**Affiliations:** Centre for Mechanochemical Cell Biology, Warwick Medical School, Gibbet Hill, Coventry CV4 7AL

## Abstract

Human Kif15 is a tetrameric kinesin-12 that contributes critically to bipolar spindle assembly in eukaryotes. Here we examine its single molecule mechanics. Under hindering loads, Kif15 steps predominantly towards microtubule plus ends, with its forestep:backstep ratio decreasing exponentially with load and stall occurring at ~6pN. Between steps, Kif15 binds stably, usually via a single head domain. By complete contrast, under assisting loads, Kif15 detaches rapidly, even in AMPPNP. Furthermore, Kif15 can autoinhibit, via an interaction requiring its C-terminus. Autoinhibited Kif15 binds microtubules nucleotide-independently, resists both hindering and assisting loads, and is further stabilized by Tpx2, which interacts with the Kif15 C-terminus. Our data reveal the mechanics of Kif15 to be extraordinarily sensitive to loading direction. When unloaded, it walks rapidly; when pulled forwards it slips and when pulled backwards it grips. We discuss the implications of this unique mechanical behaviour for the roles of Kif15 in spindle function.

## Introduction

Kinesin molecular motors play pivotal roles in the assembly and maintenance of bipolar spindles (Cross and McAinsh 2014). By transducing the energy freed by ATP hydrolysis, kinesins generate impulses of force and movement that are essential for spindle self-organisation. Kif15 (hKlp2) is a Kinesin-12 whose role in human spindles is partially redundant with that of the kinesin-5, Eg5. Eg5 is essential for spindle pole separation, but dispensable for maintaining spindle bipolarity, provided Kif15 is functional. Kif15 can support spindle assembly in the absence of Eg5, either when overexpressed or in the absence of dynein (Tanenbaum, Macurek et al. 2009, van Heesbeen, Tanenbaum et al. 2014). Eg5 and Kif15 both move towards microtubule (MT) plus ends and counteract the actions of the minus end directed motors, dynein and kinesin-14 (van Heesbeen, Tanenbaum et al. 2014). Beyond this, it is unclear whether the molecular mechanisms of KIf15 and Eg5 are similar. The functional overlap between Kif15 and Eg5 could allow cells to develop resistance to Eg5 inhibiting drugs, which are being developed as antiproliferative agents (Rath and Kozielski 2012). It is therefore important both for fundamental and practical reasons to understand the molecular mechanisms by which Kif15 motors move and exert forces on microtubules.

Several cell biological studies have suggested that the functional redundancy of Kif15 with Eg5 might reflect a similar stepping mechanism and a similar action on microtubules (Tanenbaum, Macurek et al. 2009, Vanneste, Takagi et al. 2009). By contrast, recent *in vitro* reconstitution work on the action of single Kif15 tetramers on microtubules has revealed Kif15-specific behaviours (Drechsler, McHugh et al. 2014, Drechsler and McAinsh 2016, Mann, Balchand et al. 2017). Unlike Eg5 (Valentine, Fordyce et al. 2006), Kif15 motors at zero load are highly processive, being capable of moving over 1μm along a microtubule before detaching. Walking Kif15 tetramers display a combination of paused, diffusive and motile behavior. Kif15 motors have also been shown to reduce catastrophe rates of dynamic MTs, cross-link pairs of MTs and promote assembly of parallel bundles (Drechsler and McAinsh 2016). This would be consistent with the proposed *in vivo* role for Kif15 in stabilising parallel K-fibres (Sturgill and Ohi 2013). The localisation of Kif15 to K-fibres (and interpolar MTs) is dependent upon Tpx2 (Tanenbaum, Macurek et al. 2009). Tpx2 has been shown to cause both Eg5 and Kif15 motors to bind statically to MTs *in vitro* (Drechsler, McHugh et al. 2014, Balchand, Mann et al. 2015, Mann, Balchand et al. 2017).

In contrast to this increasingly clear picture of Kif15 cell biology at low loads, little is known about Kif15 mechanics under load, either at the single molecule level or at the ensemble level in spindles. Here we aim to illuminate this problem by interrogating the mechanics of single Kif15 molecules under calibrated loads. We find that Kif15 behaves as an active molecular ratchet, walking long distances at zero load, arresting and gripping the microtubule under hindering (backwards) load, and slipping under assisting (forwards) load. This acute sensitivity to loading direction is distinctive, and shows that Kif15 motors are optimized for a different *in vivo* mode of operation to Eg5, creating the same result (a well-organised bipolar spindle) by a different organizational pathway, with both Eg5-specific and Kif15-specific spindle self-organisation likely regulated by Tpx2.

## Results

### Single Kif15 motors step processively plus endwards under load

In this study we examined a full length Kif15 construct, Kif15-FL, and a truncated construct, Kif15-1293, which lacks the C-terminal region established to bind Tpx2 (Tanenbaum, Macurek et al. 2009)(***Figure 1A***). We bound single Kif15 motors to polystyrene beads, at a dilution designed to ensure one molecule per bead (see *Methods*), and observed their behavior as they walked along MTs under load at 1mM ATP (***Figure 1B***). Kif15-FL single molecules are processive under load, but by comparison with Kinesin-1, their stepping is slower, run lengths are shorter and the motors take fewer steps at high loads.

**Figure 1.**
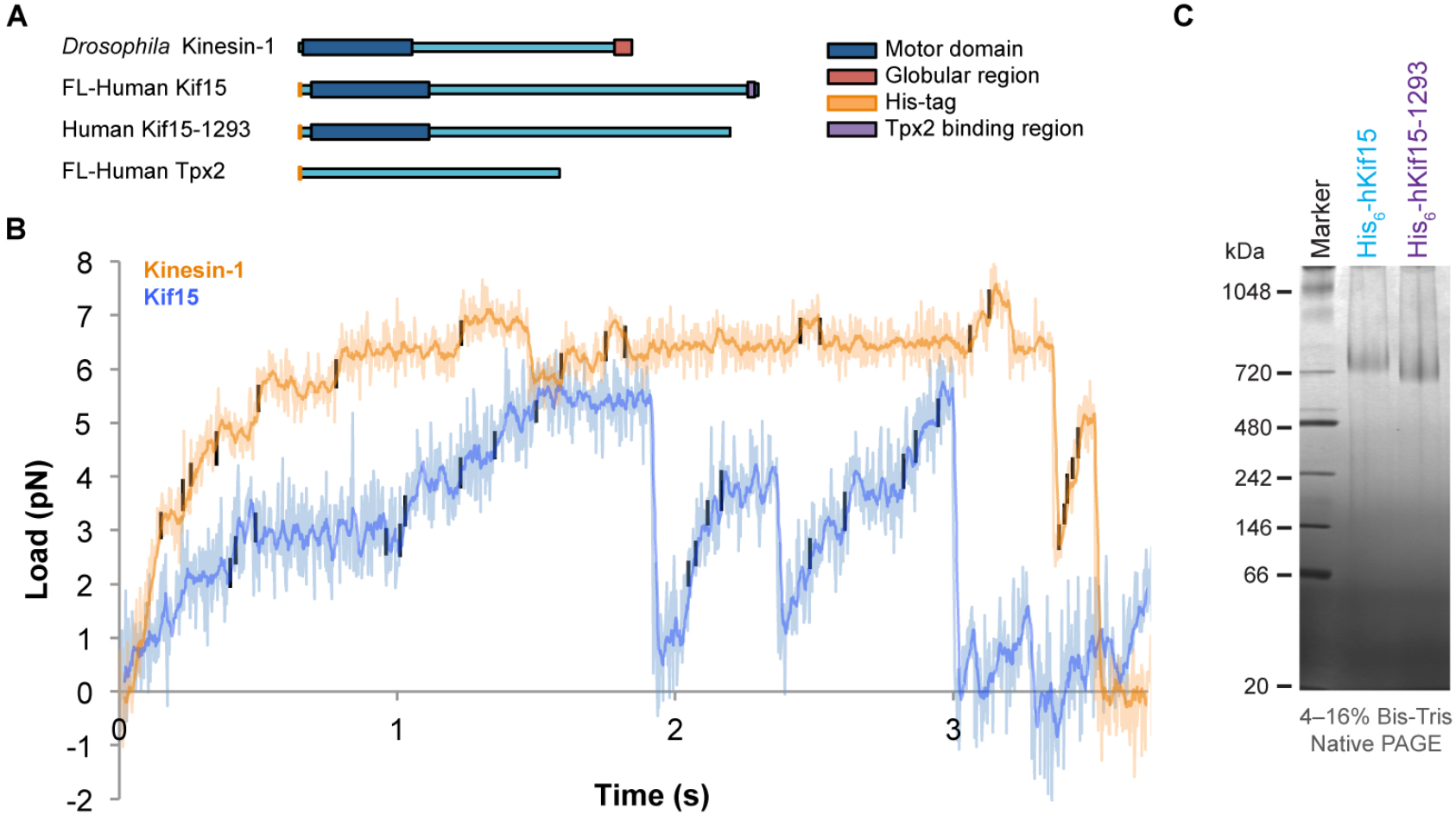
Kif15 steps towards microtubule plus ends under loads of up to 5pN. (**A**) Schematics of the 4 proteins used in these experiments; Full length *Drosophila* kinesin-1, full length hKif15 with N-terminal His-tag (Kif15-FL), C-terminally truncated hKif15-1293 lacking the terminal 95 a.a. including the Tpx2 binding region (Kif15-1293), and full length hTpx2. (**B**) Kif15-FL (blue) and Kinesin-1 (orange) stepping under load in 1mM ATP. In these example traces both the Kif15-FL and Kinesin-1 motors slow to a stall at around 6pN. Both show clear 8nm steps in both the forwards and backwards directions. Trap stiffness’s are 0.095 pN/nm and 0.081 pN/nm for FL-Kif15 and Kinesin-1 respectively. Walking Kif15 reaches lower loads and steps more slowly than walking kinesin-1. Steps were detected using a t-test based algorithm (*see Methods*) and are marked in black. (**C**) Mobility of Kif15-FL and Kif15-1293 on non-denaturing PAGE. Both constructs are tetrameric.

The run lengths of Kif15-FL and Kif15-1293 under load (***Figure S1C and S2B***) are much shorter than the run lengths observed at zero load, where frequent runs of over 1 bead diameter (560nm) are seen, and in earlier single molecule TIRF experiments (Drechsler, McHugh et al. 2014). At low trap stiffnesses, the short processive runs of steps that we observe end in detachment at low (1-3 pN) loads. As shown in ***Figures S1A and B***, at higher trap stiffness Kif15-1293 reaches higher loads before detaching, showing that progress away from the trap centre is limited by the limited run length of the motor and not by an inability to step against higher loads. By increasing the trap stiffness, the motor needs to take fewer steps to reach high loads, and correspondingly higher detachment loads are reached.

On detachment, Kif15-1293 returns either rapidly or slowly to the trap centre (***Figure S1E***) suggesting that on some occasions, the motor remains in contact with the MT even when the motor domains are no longer strongly-bound to the MT. This behavior is seen for both Kif15-FL and Kif15-1293.

At ionic strengths in the physiological range both Kif15-FL and Kif15-1293 are tetrameric (***Figure 1C***). At higher salt concentrations these tetramers dissociate into dimers (Drechsler, McHugh et al. 2014). To look for functional differences between Kif15-FL dimers and tetramers, we dissociated the tetramers by incubation at high ionic strengths before attaching them to beads and injecting into the flow cell. Exposure to high ionic strength had no effect on the measured mechanical characteristics of Kif15-FL (***Figure S2***). This suggests that the behavior we see is due to the action of just one pair of Kif15 motor domains. In support of this, steps detected in both Kif15-FL and Kif15-1293 traces using an automated step detection algorithm were of a similar amplitude to Kinesin-1 steps (***Figure S3***). If both head-pairs of the tetramer interacted with the MT, we would expect step size to vary, especially at higher loads, since in this case the load would necessarily distribute between the two pairs of engaged heads.

### Auto-inhibition of Kif15-FL requires its C-Terminal Tail

In our assays we did not see alternation of motors between long periods of paused and processive movement, as seen in single molecule TIRF assays (Drechsler, McHugh et al. 2014). Instead, motors were either stepped or adopted their tightly-bound (“locked”) state. Applying load to this locked state did not induce stepping. For Kif15-FL, only 27% of motor-bead complexes moved processively. The remainder bound statically to the MT lattice (***Figure 2A***). Upon introduction of Kif15 C-terminal targeting antibodies to the running buffer, the fraction of bead-motor complexes that moved processively was increased to over 60%, showing that the previously-observed lack of processivity is not due to permanently inactive motors in the preparation. The antibody is added after motor-bead binding and immediately before imaging and thus is unlikely to affect the bead-binding configuration of the motor. We speculate that, as with many other motors, Kif15 autoinhibition occurs via a head-tail binding interaction. In support of this idea, deletion of the terminal 95 amino acids (Kif15-1293) increased the overall motile fraction of bead-motor complexes to 84%. This fraction moved at slightly increased velocities compared to FL-Kif15 (***Figure 2A, B***). The C-terminal subdomain of the Kif15 tail is thus key to the inhibitory mechanism of Kif15. By deleting this region, or blocking it with an antibody, Kif15 auto-inhibition is relieved. Conversely, we suspect that transient autoinhibition may occur in the Kif15-FL motor, so as to reduce its average stepping rate.

**Figure 2.**
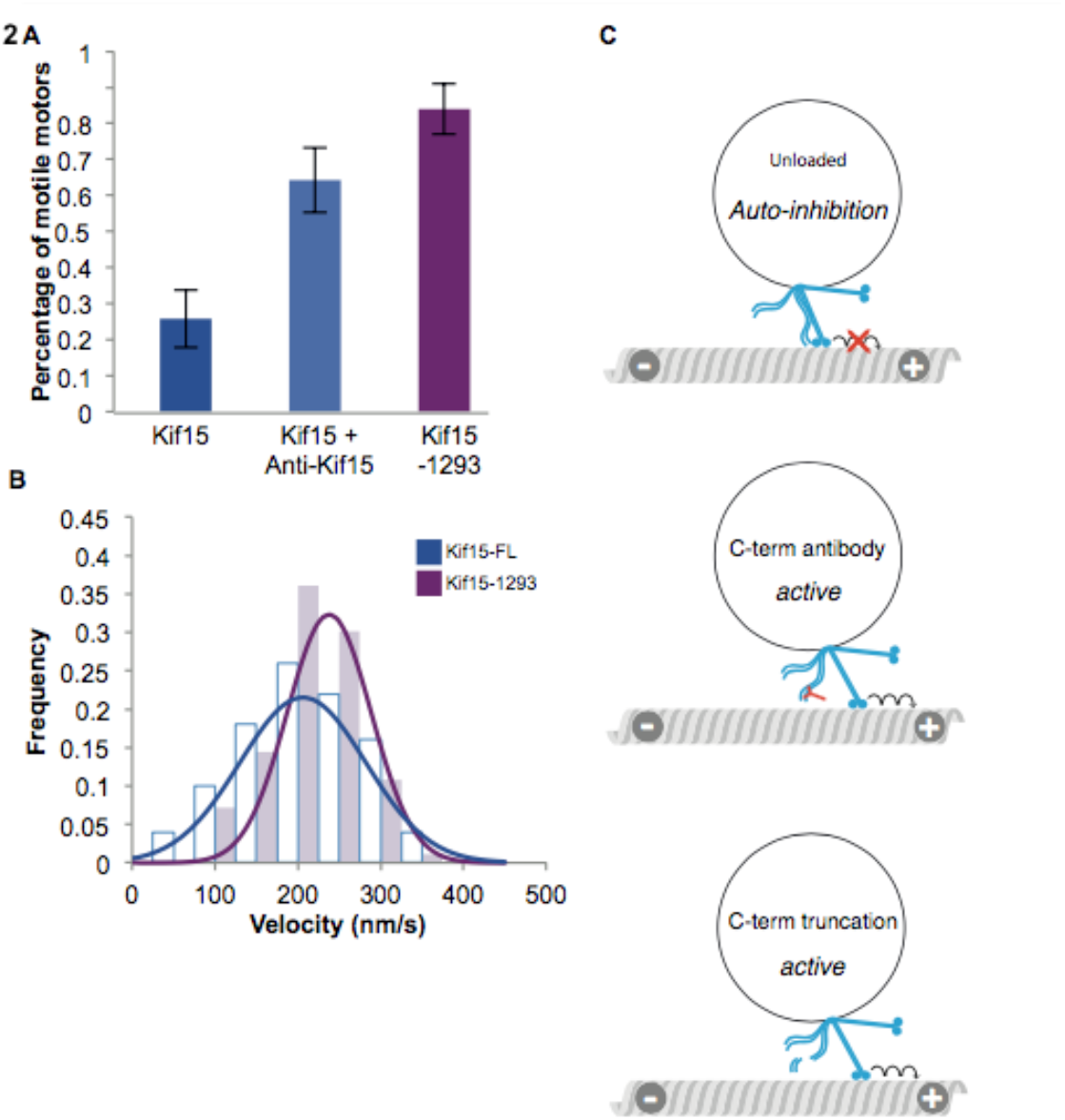
The C-terminal tail of Kif15 is able to inhibit motor stepping and this inhibition is enhanced by Tpx2. (**A**) Proportion of moving beads from those that bind the MT for Kif15-FL, Kif15-FL in the presence of 1mg/ml anti-Kif15 antibody and for Kif15-1293. Error bars show region of 95% certainty. (**B**) Unloaded velocities of Kif15-1293 motors (blue outline) and Kif15-FL motors (purple). The velocity of Kif15-1293 is 238±6nm/s, and of Kif15-FL is 206±10nm/s mean and SE. (**C**) Speculative model of Kif15 inhibition. The tail of Kif15 can bind the motor domains and the motor stops processive movement (top). In the presence of antibodies binding of the tail to motor domains is disrupted and autoinhibition is blocked (middle). For Kif15-1293 the end of the tail is removed and the tail cannot bind to and inhibit stepping by the motor domains (bottom).

Since in our experiments the attachment of the motors to beads relies on nonspecific binding interactions, it is likely that the motors are attached in multiple different ways. We characterized this variation to some extent by measuring the tether length of the bead-MT connection (***Figure S4***). For Kif15-FL this was more variable than for Kinesin-1 indicating that Kif15 binds to the beads in a variety of configurations (***Figure 2C***).

### Measured force-velocity relationship for Kif15

Although the run lengths of Kif15-FL and Kif15-1293 are much reduced under backwards load, most molecules nonetheless take several steps at the trap stiffnesses we used (***Figure S1A***). Kif15-1293 does not auto-inhibit and steps consistently under load, allowing us to collect stepping data over a range of loads. For Kif15-1293 and for Kinesin-1, which served as our positive control, both forwards and backwards steps were seen. ***Figure 3A*** shows the average dwell times (the time interval preceding each step) under load in 1pN bins for both motors. We pooled the measured dwell times for both forwards and backwards steps. These data show clearly that Kif15-1293 displays longer dwell times than Kinesin-1 at all loads, corresponding to a slower stepping rate.

**Figure 3.**
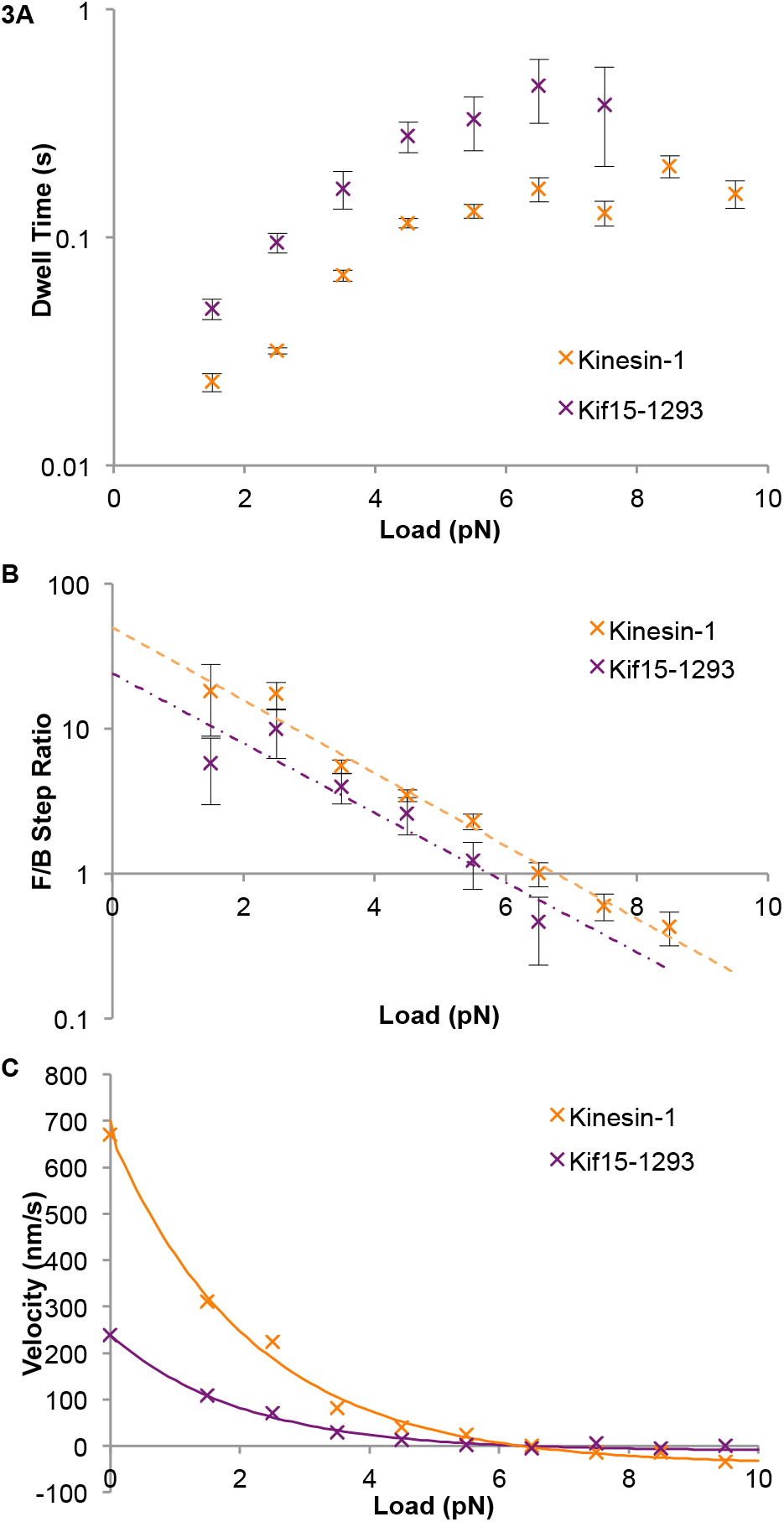
Kif15 steps slower than Kinesin-1 at all loads and has a similar stall force. (**A**) Dwell time distributions for Kinesin-1 and Kif15-1293 in 1mM ATP. A dwell is the time that the motor waits between steps. Both motors show an increase in dwell time at higher loads. Above 6.5pN, Kinesin-1 dwell times plateau. (**B**) Forwards/Backwards stepping ratios for 1pN binned data for Kinesin-1 and Kif15-1293. Fitted lines are exponentials, fit using least squares. Stall forces (when the number of forwards and backwards steps are equal) are 5.6±0.76 pN for Kif15-1293 and 6.76±0.76 pN for Kinesin-1. Errors are calculated from the standard error in the slopes and intercepts of the least squares line fits. (**C**) Dependence of velocity on load for Kif15-1293 (purple) and Kinesin-1 (orange). Velocity is calculated from dwell times and Forwards/Backwards stepping ratio at each load, assuming a step size of 8.1nm for both motors. Zero load data points are from unloaded bead velocities. Kif15-1293 and Kinesin-1 distributions are fit with exponentials, for parameters see Supplementary Table 1.

By counting the number of forwards and backwards steps made by a motor at each load it is possible to determine its stall force, the backwards load at which the motor makes an equal number of forwards and backwards steps and so makes no net progress. ***Figure 3B*** plots the ratio {number of forwards steps}:{number of back steps} versus load, using 1pN bins and comparing Kinesin-1 with Kif15-1293. By fitting exponentials, we determined the stall forces of the motors by extrapolation.

We measure a stall force for Kinesin-1 of 6.8±0.7pN, in quantitative agreement with previous data (Carter and Cross 2005). For Kif15-1293, we measure a stall force of 5.8±0.7pN. The load-dependence of the dwell times for Kif15-1293 and Kinesin-1 shows a similar exponential factor, but the stall force for Kif15-1293 is slightly lower than that of Kinesin-1. Above stall force (super–stall) both Kif15-1293 and Kinesin-1 motors are capable of making several successive backwards steps. Published data for Eg5 dimers predicts a similar high stall force of at least 7pN (Valentine, Fordyce et al. 2006) although both dimeric and tetrameric Eg5 are reported to dissociate from the MT rather than slow their velocity (Valentine, Fordyce et al. 2006, Korneev, Lakamper et al. 2007).

### Unbinding of Kif15-1293-AMPPNP under load depends acutely on loading direction

Even modest hindering (backwards) loads substantially reduce the processivity of Kif15 motors. To examine the detailed load-dependence of Kif15 detachment in different nucleotide states, we performed force-ramp experiments. In these experiments, which were similar to those of Uemura et al., 2002, the stage was moved at a constant rate, sliding a MT past a trapped, bead-attached motor that remains engaged with the MT (***Figure 4A***). Slowly transporting the MT by moving the microscope stage then steadily increases the load on the motor-MT connection. Eventually the motor unbinds from the MT and the bead-attached motor returns to the trap centre. Example bead position recordings are shown in ***Figure 4B-F***. By recording the loads at which the motor-bead complex detached from the MT under assisting (plus-end directed) or hindering (minus-end directed) loads, we were able to quantitate the dependence of the unbinding rate on load, for plus and minus end directed loads, for both Kif15-1293 and Kinesin-1 and in both stepping and non-stepping conditions.

**Figure 4.**
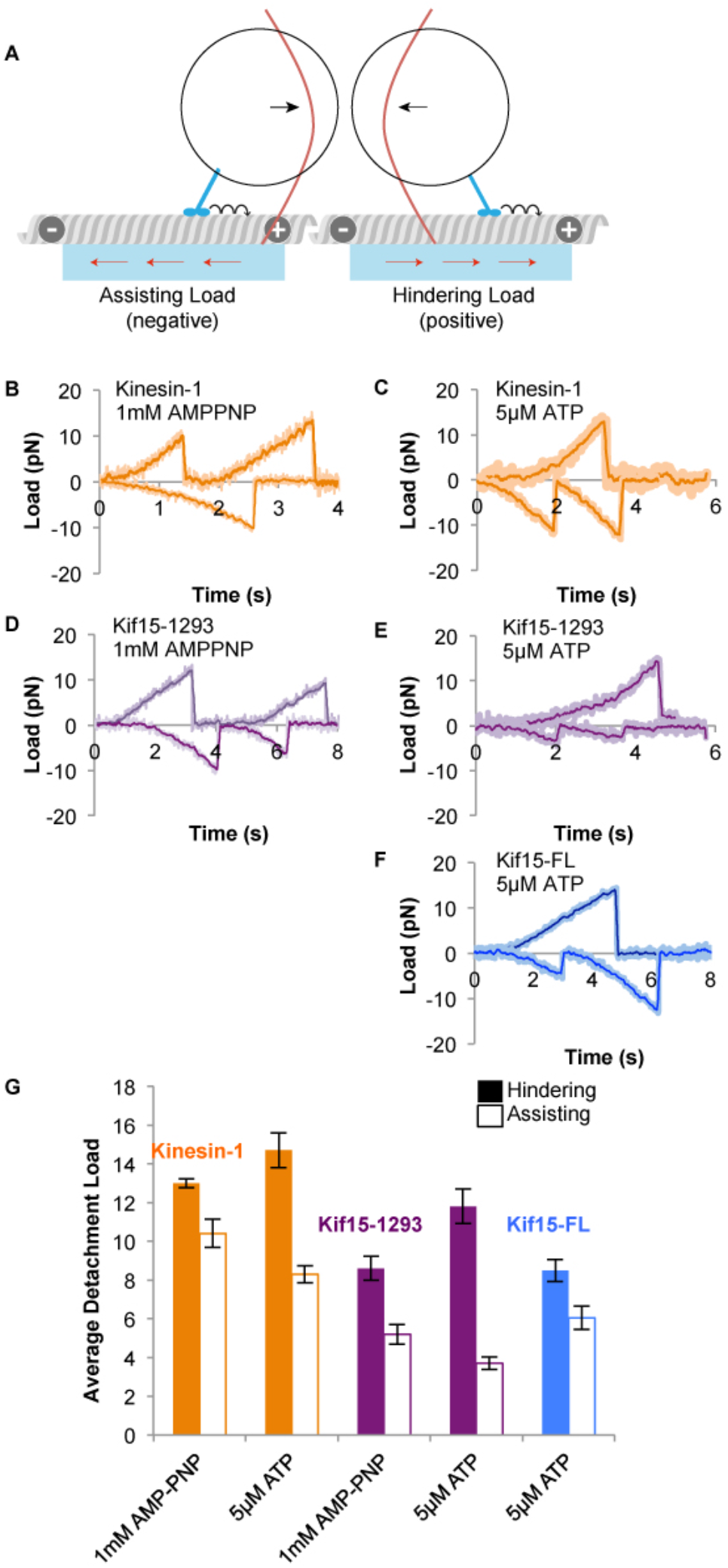
Kif15 unbinding from MTs is very sensitive to loading direction. (**A**) Experimental set up for applying assisting (negative) and hindering (positive) loads. With the motor bound to the MT, the stage is moved at a defined rate (red arrows) so as to drag the MT relative to the bead-attached motor in the trap centre. The bead-attached motor is thereby steadily translated out of the trap centre and experiences a steadily-increasing restoring force. The position of the bead is tracked continuously and the position and load at which the motor detaches from the MT is recorded. (**B,C**) Example traces for kinesin-1 in the presence of 1mM AMP-PNP (**B**) and 5μM ATP (**C**). Positive values indicate hindering loads whilst negative values represent assisting loads. (**D & E**) Example traces for Kif15-1293 in the presence of 1mM AMP-PNP (**D**) and 5μM ATP (**E**). Positive values indicate hindering loads whilst negative values represent assisting loads. (**F**) Example traces for Kif15-FL in the presence of 5μM ATP. (**G**) Comparison of average detachment loads in 5μM ATP or 1mM AMPPNP under hindering (filled bars) and assisting (unfilled bars) loads. Kinesin-1 (orange), Kif15-1293 (purple) and FL-Kif15 (blue). Detachments under hindering loads occur at higher forces than those under assisting loads for all motors and nucleotide conditions.

Results for Kinesin-1 in the presence of the slowly hydrolysable ATP analog 1mM AMP-PNP closely resemble those of Uemura et al., 2002. In these conditions, AMP-PNP is saturating but hydrolysis does not occur and stepping rates of Kinesin-1 are not measurable. In 1mM AMP-PNP, Kinesin-1 unbinds more rapidly under assisting loads. At the pulling-rates we used (***Figure S5***), unbinding under minus end directed (hindering) load requires 25% more load than unbinding under assisting loads (***Figure 4B and G***). As noted by Uemura et al., unbinding loads of kinesin-1 in AMPPNP show a bimodal distribution, thought to represent motors unbinding from either a single or double head bound state (***Figure 5***).

**Figure 5.**
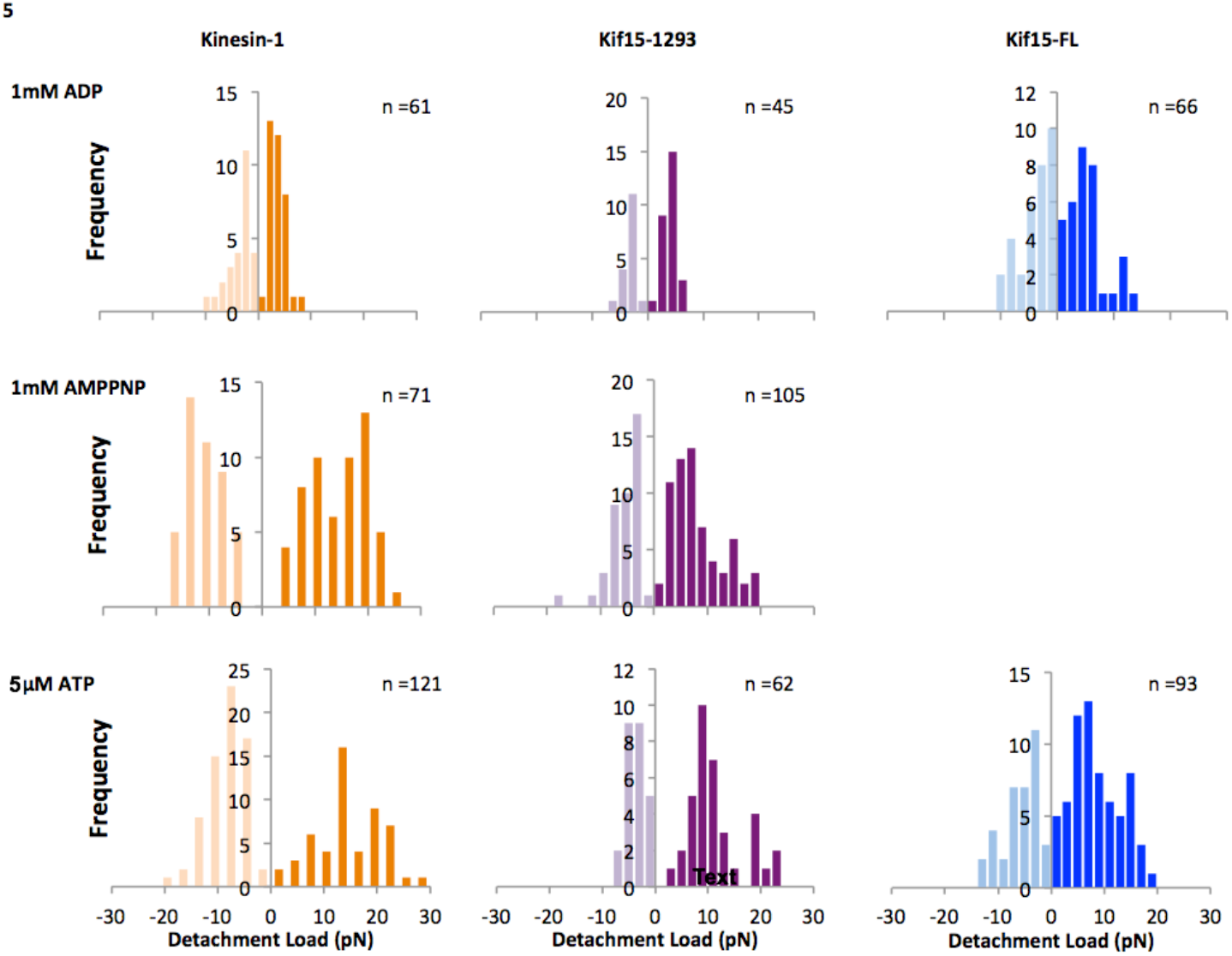
Detachment load distributions for kinesin-1, Kif15-1293 and Kif15-FL under hindering (positive) and assisting (negative) loads. Loading rates are 3.6±1.1 pN/s for ADP data and are shown in Figure S5 for AMP-PNP and ATP data.

In the same conditions, Kif15-1293 requires 67% more load to detach the motor from the MT under minus end directed loads than under plus end directed loads (***Figure 4C and G***). The mean unbinding forces for Kif15-1293 (8.6pN for hindering loads, 5.2pN for assisting loads) are lower than those for Kinesin-1 (respectively 13.0pN and 10.4pN). This reduced stability is consistent with the 12.2s residence time previously reported for unloaded C-terminal truncated (1-700aa) Kif15 in 1mM AMP-PNP (Sturgill, Das et al. 2014) compared to ~100s for Kinesin-1 (Hancock and Howard 1999). From our results in 1mM AMP-PNP it appears that the AMP-PNP state of Kif15-1293 is less stably attached, and its detachment more sensitive to loading direction, than that of Kinesin-1.

The distribution of unbinding loads for Kif15-1293 in AMP-PNP under hindering loads is also asymmetrical, with a lobe of the distribution exhibiting roughly double the detachment force compared to the majority (***Figure 5***). This high force component is less prominent for Kif15 than for Kinesin-1, suggesting that in the presence of AMP-PNP Kif15 spends more time bound to the MT by a single motor head than does Kinesin-1. Measured unbinding loads for Kif15-1293 under assisting load clustered in a single peak centred on 5.2pN.

### Unbinding of Kif15-1293 during ATP-driven stepping also depends steeply on loading direction

We also investigated the unbinding of Kif15 under load during ATP-driven processive stepping. The format of this experiment is very similar to the AMP-PNP force-ramp experiment, but at 5μM ATP, motors will continue to undergo ATP hydrolysis and make steps along the MT, albeit at a much-reduced pace. At 5μM ATP the unloaded velocity of Kinesin-1 coated beads is approximately 50 nm/s or about 6 steps/s. In these conditions, Kinesin-1 unloading asymmetry becomes more pronounced, with 76% more force required to unbind motors using hindering loads compared to assisting loads (***Figure 4C and G***).

Under assisting loads the stepping rate of Kinesin-1 remains essentially unchanged (Carter and Cross 2005) whilst under hindering loads the stepping rate slows as load is applied. It has recently been found that for Kinesin-1, the run length under assisting loads is much shorter than that under hindering loads (Clancy, Behnke-Parks et al. 2011, Andreasson, Milic et al. 2015). This suggests that assisting loads disrupt the coordination of kinesin-1 stepping.

In 5μM ATP, unloaded Kif15-1293 moves along MTs at a velocity of approximately 36nm/s, corresponding to 4.5 steps/s. In these conditions the asymmetry seen for Kif15-1293 unbinding is dramatic, with 3-fold more hindering load than assisting load needed to detach Kif15-1293 (***Figures 4E and 5***). Compared to Kinesin-1, this asymmetry of the unbinding load for Kif15-1293 is due to both an increase in detachment force under hindering loads and a decrease in detachment force under assisting loads (***Figure 4G***). Under hindering loads this might be caused by increased time spent in the tighter-binding ADP.Pi and/or nucleotide-free states. As for Kinesin-1, we suspect that assisting load will cause Kif15-1293 to lose coordination, leading to early detachment. We predict that run lengths of uninhibited Kif15 under load will therefore be very heavily dependent on loading direction; this is very different to the behavior of Eg5, which shows little loading direction-dependent run length asymmetry (Valentine and Block 2009).

### Unbinding of Kif15-FL shows reduced dependence on loading direction

At 5μM ATP, Kif15-FL steps unmeasurably slowly. Potentially, under these conditions the tail of Kif15-FL might inhibit stepping by interacting with the head domains (see *Discussion*). The average unbinding loads of Kif15-FL are similar to those of Kif15-1293 in 1mM AMP-PNP (***Figure 4G***). The distribution of Kif15-FL unbinding loads in 5μM ATP shows more detachments occurring at high loads for both assisting and hindering forces. We assign these higher load detachments to the auto-inhibited (“locked”) state of Kif15-FL (see *Discussion*).

In 1mM ADP, Kif15-FL displays a small subset of detachments occurring at higher load, not seen in either the Kif15-1293 or Kinesin-1 data sets (***Figure 5***). As in the 5μM ATP conditions, we ascribe these higher-load detachments - which are dependent on the full-length tail - to a state with increased stability of motor-microtubule binding. This tail-mediated stabilization operates under both assisting and hindering loads. As with all kinesins, ADP biases Kif15-FL towards weak binding, but even in 1mM ADP Kif15-FL retains its innate directional asymmetry so that on average detachments occur at 5.2±0.5pN and 4.0±0.5pN for hindering and assisting loads respectively.

### Kif15 auto-inhibition is stabilized by Tpx2

Tpx2 is a binding partner for Kif15, implicated in the in vivo loading of Kif15 to MTs and known to inhibit Kif15 motor activity. Injection of 18nM dimeric Tpx2 into the optical trapping assay chamber enhances the dwell time of unloaded Kif15-FL motors and also greatly increases their resistance to load (Drechsler, McHugh et al. 2014). In the unbinding load experiments described above (***Figure 4A***), Kif15-FL in the presence of Tpx2 remained MT-bound at loads of over 35pN, the limit of our optical trap. This was true under both hindering and assisting loads and argues strongly that full-length Kif15 functions as a stable MT crosslinker in the presence of Tpx2. The interaction of Kif15-FL with Tpx2 occurs via its C-terminal tail domain. Kif15-1293, which is missing this domain, is relatively unaffected by the presence of Tpx2 when compared to Kif15-FL (***Figure 6A and B***).

**Figure 6.**
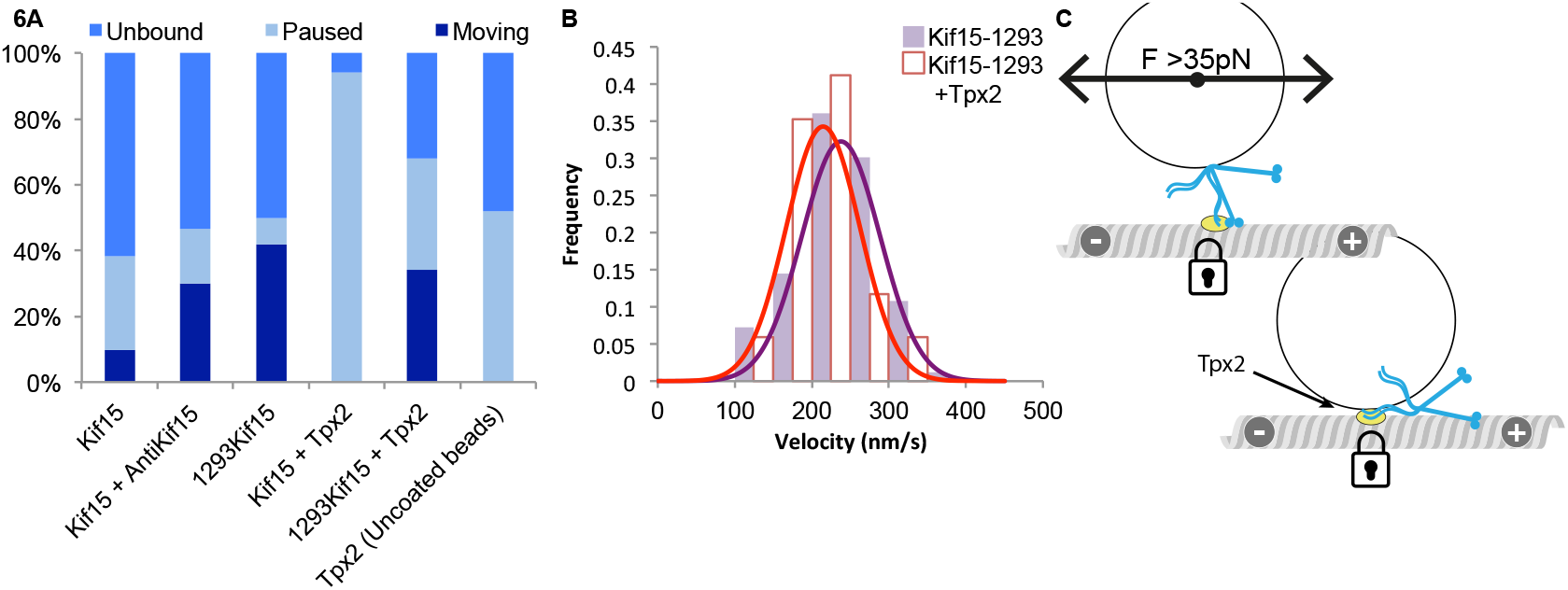
Effects of Tpx2 on Kif15 mechanics. (**A**) Fraction of bead-motor complexes assayed that were unbound, bound but paused, or bound and moving, after Kif15, Kif15-1293 or control (uncoated) beads were introduced to ≥2 MTs, for at least 30s, in the presence and absence of 18nM dimeric Tpx2. (**B**) Unloaded velocities of Kif15-1293 motors in the presence of 18nM dimeric Tpx2 (red outline) and in the absence of Tpx2 (purple). The velocity with Tpx2 is 215±12nm/s without Tpx2, 238±6nm/s, mean and SE. (**C**) Schematic of Tpx2 effect on Kif15 coated beads. Interaction of the tail with the motor domains is not necessary for Tpx2 interaction. When Tpx2 is bound to the tail of Kif15 it binds the MT tightly and will not release under loads of more than 35pN.

Under the same conditions Kif15-FL is almost always inhibited by Tpx2 (***Figure 6A***), indicating that bead-bound Kif15-FL is in a configuration in which the C-terminal part of its tail domain is always available for binding Tpx2. Since only a subset of Kif15-FL motor-bead complexes that bind the microtubule auto-inhibit, yet all are arrested by Tpx2 binding, it seems likely that Tpx2 binding occurs via an independent mechanism where binding of Tpx2 to the motor C-terminus is all that is required to arrest progress, independent of tail-mediated auto-inhibition (***Figure 6C***). It has previously been shown that Kif15-FL and Tpx2 do not bind tightly to one another in solution (Drechsler, McHugh et al. 2014), suggesting that Tpx2 binds only once Kif15-FL is attached to the MT.

## Discussion

### The mechanics of Kif15 are distinct from those of kinesin-1 and Eg5

Our data establish that single Kif15 motors have a distinct mechanochemical signature, clearly different from Kinesin-1 or Kinesin-5. Kif15 shows some functional overlap with kinesin-5 (Eg5) and at the single molecule level, Kif15 indeed shares some mechanical characteristics with Eg5. Like Eg5, Kif15 under load has a short run length and tends to detach from the MT before reaching stall. However, compared to Kif15, the mechanics of Eg5 are comparatively insensitive to the loading direction (Valentine and Block 2009). Kif15 also shows some similarities to Kinesin-1, such as its long, unloaded run length and the exponential relationship of its forestep-backstep ratio to load. However, for Kif15, the effect of loading direction on MT binding is substantially more asymmetrical than for kinesin-1, so that single Kif15 molecules detach rapidly as soon as they experience even a modest component of assisting load. These distinctive single molecule mechanics predict that in the spindle, Kif15 will function as an active ratchet, walking rapidly towards plus ends at zero load, gripping and / or stepping slowly under hindering load, and releasing quickly under assisting load (***Figure 7A***).

**Figure 7.**
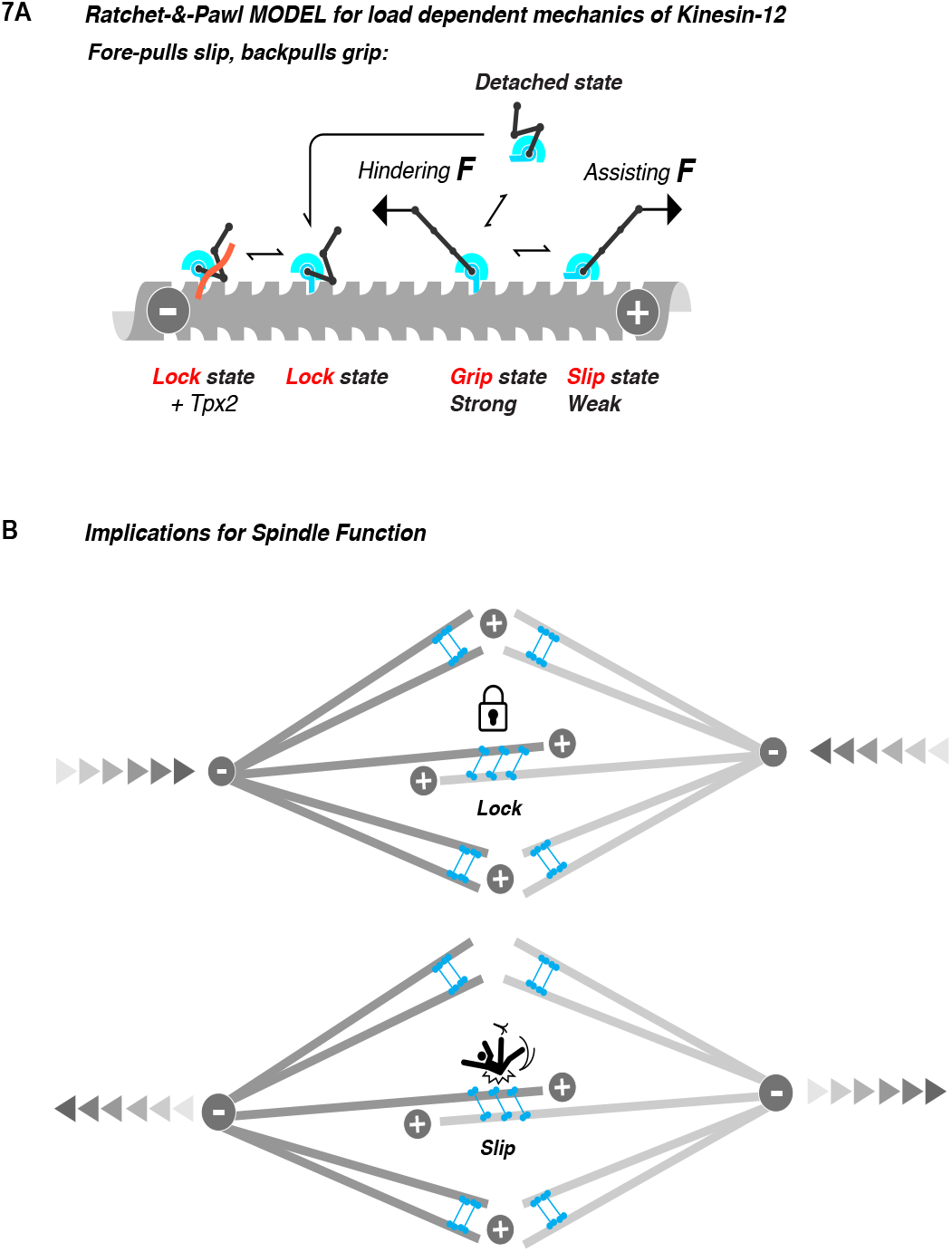
Ratchet and Pawl model for load-dependent mechanics of Kif15. Kif15 motor is represented as a ratchet-and-pawl (blue structures) that slips (slip state) when pulled towards plus ends (assisting load) but grips (grip state) when pulled in the opposite direction (hindering load). The ratchet-and-pawl structure can also adopt a lock state, in which the C-terminus of the tail interacts with the ratchet and pawl and causes it to bind the microtubule stably and nucleotide-independently. This state can be further stabilised by Tpx2. We hypothesize that the lock state can only be formed on binding of the detached state to the MT. (**A**) Kif15 can bind to microtubules in either a motile state or a lock state. Application of minus end directed (hindering) load to the motile state slows motor stepping causing it to ‘grip’ the MT. Application of assisting load causes the motile state to easily ‘slip’ along the microtubule lattice with very little resistance. The lock state of Kif15 is also sensitive to loading direction but to a lesser extent than the motile state. The presence of Tpx2 causes the motor to lock to the MT even more strongly. (**B**) For spindles under compression, Kif15 in the overlap region of the spindle will tend to lock the sliding of MTs in the central MT overlap zone (top). In contrast, for spindles subject to expansile forces Kif15 in the overlap region will easily slip, providing little friction.

The previously controversial question of the oligomerisation state of Kif15 under physiological conditions now seems settled, as motors within mitotic or interphase extracts were confirmed to be tetrameric by single molecule imaging (Mann, Balchand et al. 2017). However it remains to be established whether the functional dimer-of-dimers of Kif15 is stabilised by parallel or antiparallel interactions, or indeed a mixture of the two, as in the *Drosophila* kinesin-5 tetramer (Scholey, Nithianantham et al. 2014). In our hands, the mechanics of Kif15 under dimer conditions (high salt buffer) are indistinguishable from those under tetramer conditions (low salt buffer). We take this to mean that the functional unit in our optical trapping experiments is the twin heads of a tethered dimer. Accordingly, we focus here on *functional* bipolarity, implying only that the two head-pairs in each functional tetramer can engage two different microtubules.

Against this background of functional bipolarity, it is striking that recent reconstitution experiments show that the two ends of Kif15 tetramers can display either equivalent or non-equivalent velocities on attached microtubules depending on microtubule geometry (Drechsler and McAinsh 2016). By showing here that Kif15 motor domains can convert between two different MT binding modes (active and locked) with different mechanical characteristics, with detachment of the active mode acutely dependent on loading direction, our findings suggest a possible origin for this otherwise puzzling behavior (***Figure 7***).

### Kif15 dwells between steps in a single-head-attached state

Our force-ramp experiments identify two different attached populations of Kif15 single molecules in AMP-PNP (***Figure 5***), which we attribute, by analogy to kinesin-1 (Uemura, Kawaguchi et al. 2002), to 1-head-attached and 2-head-attached states. Force-ramp experiments in ATP have not previously been reported for any kinesin. Our experiments in ATP show two classes of attached state, for both kinesin-1 and Kif15, but with the 1-head-attached state dominating. Thus, Kif15 dwells between steps predominantly in the state interpreted as a 1-head attached state. This suggests that the acute load-direction sensitivity of Kif15 is an intrinsic property of each Kif15 head. The single headed wait state and acutely asymmetric load-sensitivity of Kinesin-1 and Kif15 stand in sharp contrast to the mechanics of Eg5, for which recent evidence suggests that relatively load-insensitive Eg5 dimers attach to the microtubule via both motor domains for much of their mechanochemical cycle (Chen, Mickolajczyk et al. 2016).

### Kif15 can autoinhibit. Straining the tether may block autoinhibition

We find that in addition to its processive stepping mode, Kif15 can bind statically, in a stably-bound refractory (lock) state. In our hands, this state was never seen to arise during a processive run under load, although when unloaded the full-length motor does show a decreased average velocity compared to Kif15-1293. Rather, a typical pattern was that following a processive run in the laser trap, the motor would detach, and then re-attach either in an active state that underwent a further processive run, or in an inactive state that bound the MT nucleotide-independently and was resistant to both hindering and assisting loads. Our data thus suggest that entry to the refractory, autoinhibited state of Kif15 may only occur at zero load. Certainly, we never observed a train of processive steps under load that was terminated by locking of the motor to its track – processive runs always ended in a detachment. We therefore tentatively hypothesise that stretching out the Kif15 tether blocks the autoinhibition reaction. ***Figure 7*** and legend summarise these ideas as a mechanical cartoon.

### Tpx2 super-stabilises the Kif15-MT interaction

The auto-inhibited, locked state Kif15 can be further stabilized by interaction with Tpx2. In the Tpx2-stabilised lock-state, detachment from the MT requires loads >35pN in any direction. The same C-terminal 95aa region of the Kif15 tail that is required for auto-inhibition includes the Tpx2 binding region. Deletion of this region dramatically reduces the effect of Tpx2 on Kif15 (***Figure 6A***), so that velocities of motile Kif15-1293 motors are essentially unaffected by the presence of Tpx2 (***Figure 6B***). Tpx2 has previously been shown to cause Kif15 to pause on the MT lattice for extended periods of time (Drechsler, McHugh et al. 2014). Both Kif15 and Eg5 can undergo 1-D diffusion on MTs and Tpx2 also causes Eg5 to bind statically to MTs. For Kif15 it remains unclear how lattice diffusion arises. Nonetheless, Eg5 and Kif15 have it in common that interaction with Tpx2 clearly switches off the ability to diffuse, instead stabilizing a statically-bound lock-state.

### Kif15 shows reduced processivity under load

Kif15 is highly processive at zero load, but even modest hindering loads dramatically reduce processivity. In principle, the acute load-direction dependency of the Kif15 detachment rate might be expected to increase processivity, in that forwards load on the trail head enhances its detachment, whilst backwards load on the leading head suppresses its detachment. Since this clearly does not happen, we speculate that since even the “strong” nucleotide states of Kif15 detach rapidly under modest forwards loads, the trail head of Kif15 is not protected from premature detachment in the same way that the trail head of kinesin-1 is protected. Notwithstanding, it is clear that once committed to step, the fore-back bias of Kif15 is both highly effective and highly load-dependent, so that Kif15 has a stall force approaching that of kinesin-1.

### How Kif15 may operate in spindles

There is now considerable evidence that Kif15 in mitosis binds both in the overlap zone, and to K-fibres, where it counteracts inwards forces by crosslinking and lending additional stability to K-fibres. Our single molecule mechanics data support a model in which Kif15 shifts between these two roles, likely regulated by the binding of Tpx2 to the C-terminus of the tail.

As we have shown, the C-terminal tail of Kif15 also mediates an autoinhibition reaction that creates a lock state that binds tightly and nucleotide-independently to MTs. C-terminal inhibition, stabilized by Tpx2, may allow Kif15 to stably crosslink MTs in some situations. One scenario is that Kif15 motors could land in Tpx2-enriched areas of the spindle (close to the poles) and then lock to the MT, creating strong and localised crosslinks. Kif15 at MT plus ends has recently been shown to suppress MT catastrophe, again consistent with a role in K-fiber stabilization.

How does Kif15 replace the function of kinesin-5? Our evidence points to Kif15 supplementing or substituting Eg5 activity using a related but different mechanism. The salient difference between the Eg5 and Kif15 is the asymmetrical dependence of Kif15 detachment rate on loading direction. Eg5 is directionally insensitive, enabling it to act as a brake to slow the sliding apart of overlapping MTs (Shimamoto, Forth et al. 2015). In cells, Eg5 uses this mechanism to limit the rate of spindle elongation (sliding of overlapping MTs) during anaphase, a process in which Kif15 is not involved (Saunders, Powers et al. 2007).

Both Kif15 and Eg5 bind Tpx2, suggesting the possibility of linked regulation and complex emergent behavior. When interacting with Tpx2, Kif15 will function as a highly effective static crosslinker to resist sliding driven by other forces. In this situation, fluxing of the K-fibres may drive spindle pole separation. Importantly, we have shown that under forwards load (arising for example if the spindle poles are driven apart by other forces) and in the absence of Tpx2, Kif15 will detach readily. Thus, in situations where attached Kif15 motor domains are pulled in the progress direction (towards MT plus ends), they will slip readily, ensuring minimal drag.

Under hindering load, Kif15 engages relatively stably, but we have shown that its run length, previously thought to be very long, in fact shortens dramatically under even modest loads, becoming similar to that of Eg5 and potentially enabling easier teamwork between the two motors. Exploring this will require further work.

In conclusion, our data show that the single molecule mechanics of Kif15 allow it to function as an active mechanical ratchet that walks rapidly towards plus ends when unloaded, slows down and become less processive under hindering load, slips easily under forwards load, and transitions stochastically to a tight-binding, autoinhibited state at zero load. Kif15’s ability to slip under assisting load will allow it to function effectively in teams involving other plus end directed steppers. Its ability to transition to a Tpx2-stabilised autoinhibited state allows it to convert from active to static, load-independent binding. Further investigation into the spatial regulation of Kif15 by Tpx2, Ki67, Aurora A or other mitotic regulators in cells will be needed to understand the incontext global contribution of Kif15 to spindle formation and maintenance.

## Acknowledgements

R. A. Cross is supported by a Wellcome Trust Senior Investigator Award [grant number 103895/Z/14/Z]. A.D. McAinsh is supported by a Wellcome Trust Senior Investigator Award [grant number 106151/Z/14/Z] and a Royal Society Wolfson Research Merit Award [grant number WM150020]. T. McHugh was funded by an EPSRC DTG doctoral award [grant number 1090393].

## SUPPLEMENTARY FIGURES

**Figure S1.**
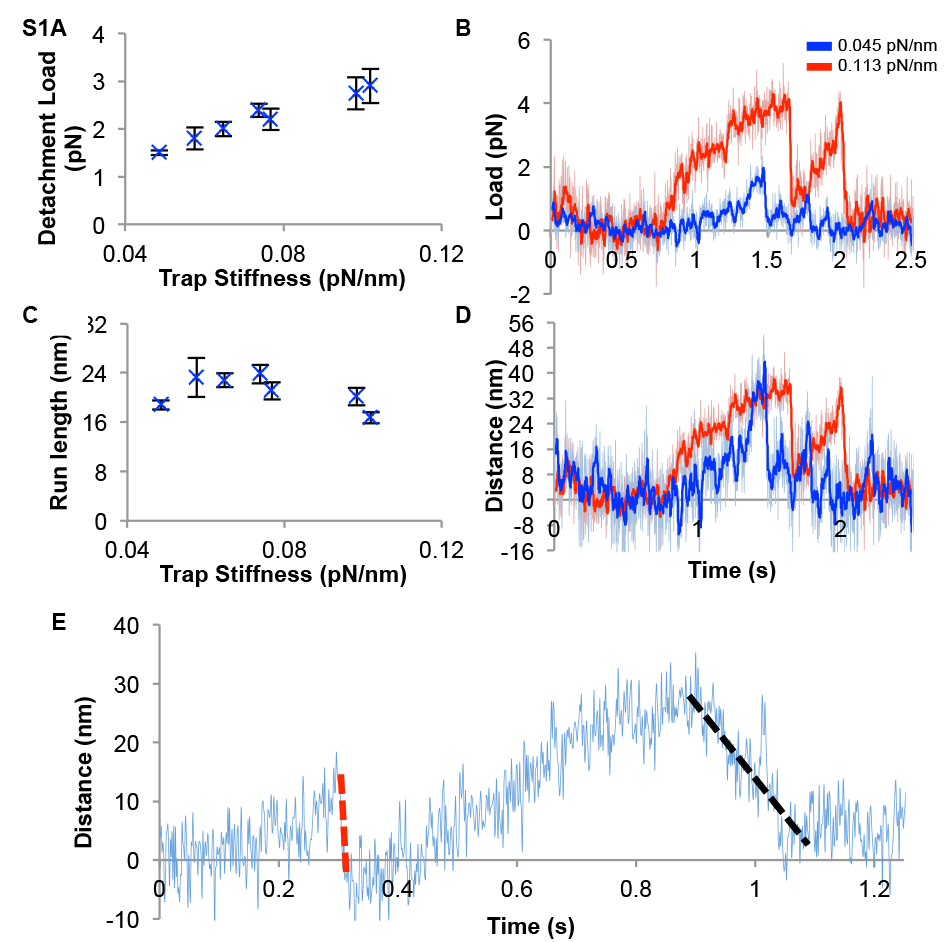
Kif15 has a very short run length under load. Average detachment loads (A) and distances (C) for a series of traces from the same Kif15-1293 motor measured at different trap stiffness’s. The average detachment load increases with increasing trap stiffness. (B & D) show example Kif15-1293 traces for the same bead taken at trap stiffnesses of 0.045pN/nm (blue) and 0.113pN/nm (red), with the two traces scaled together according to load (B) or travel distance (D). (E) Example trace showing both a fast (red) and slow (blue) return to trap centre.

**Figure S2.**
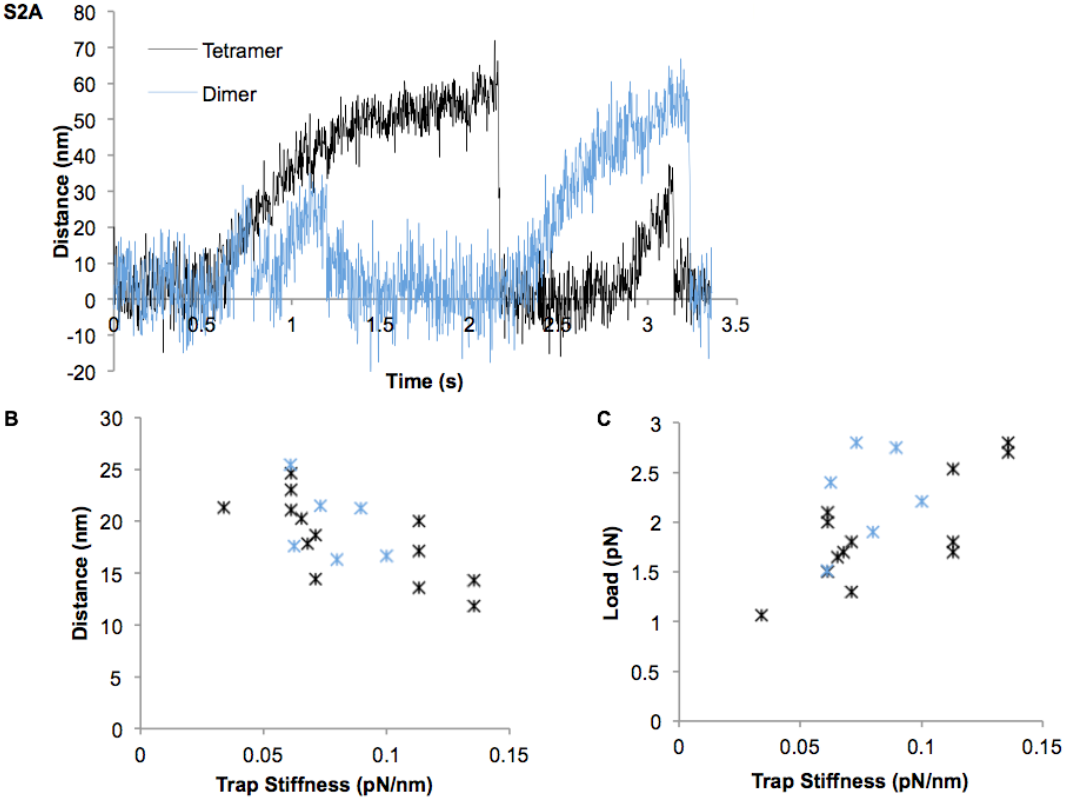
(A) Example traces for FL-Kif15 in dimer-favouring (blue, high-salt) and tetramer-favouring (black, low-salt) bead incubations. Trap Stiffness’s of 0.07 and 0.09 pN/nm were used for high and low ionic strengths respectively. (B & C) Mean detachment loads (B) and distances (C) for motors after dimer-favouring (blue) and tetramer-favouring (black) bead incubations under varying trap stiffness’s.

**Figure S3.**
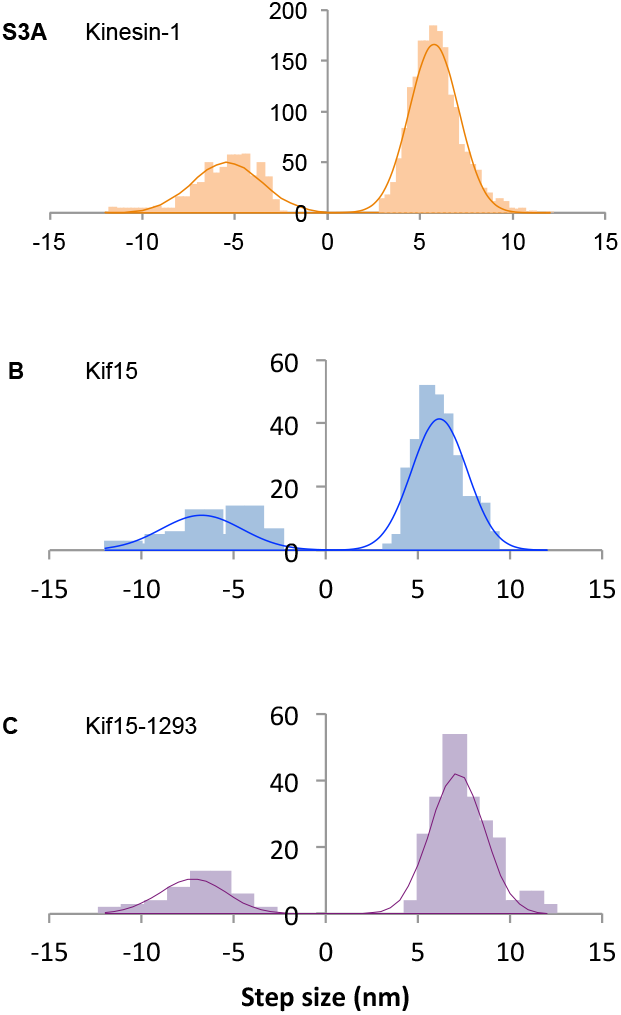
Step Size distributions for (A) Kinesin-1 (B) FL-Kif15 and (C) Kif15-1293. Normal distribution curves have been fit to the forward (plus end directed) and backward (minus end directed) step distributions. The number of steps detected and the parameters for these curves are given in Table S3.

**Figure S4.**
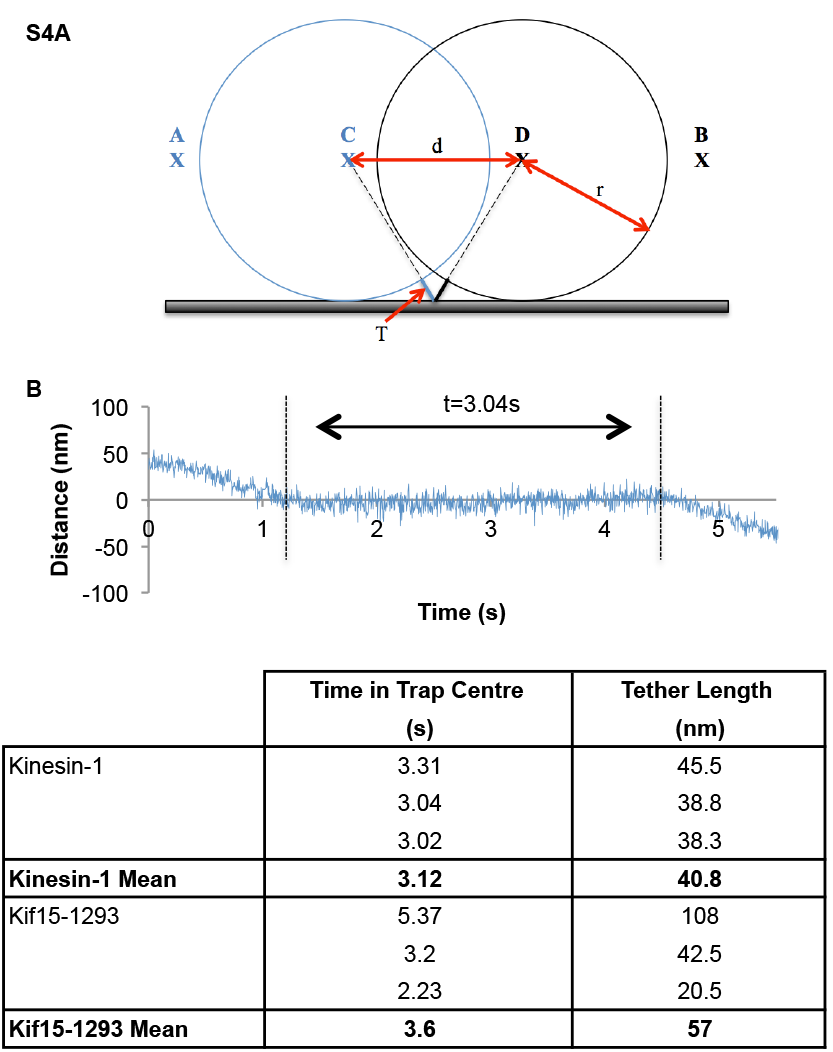
Gauging the length of the motor tail that projecting from the beads. By translating the optical trap, the motor is moved from backwards tension, through a slack state and into forwards tension. (A) The laser focus is moved from point A to point B at a constant rate of 100nm/s. For a motor of tether length T a bead of radius r will move distance d, whilst the laser focus is within the range of C and D the bead will not appear displaced from the trap centre.(B) Example trace for Kinesin-1, time in the trap centre is measured by eye to be 3.04s. Inferred tether lengths are given in Table S2.

**Figure S5.**
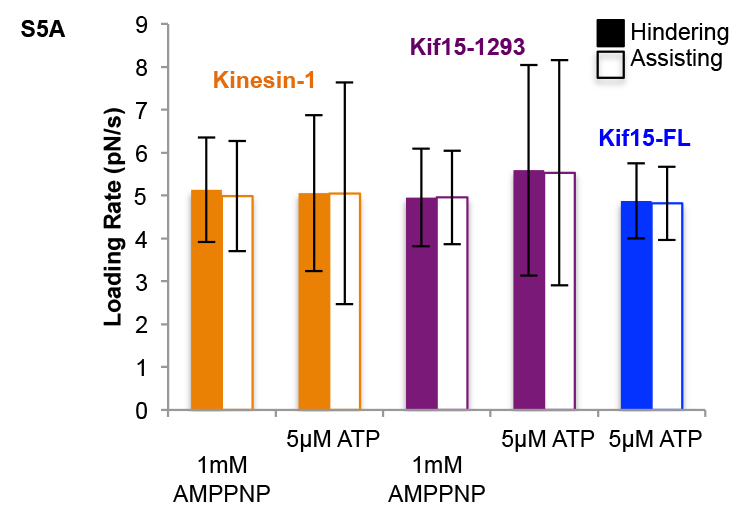
Loading rates for the data sets shown in Figure 4, mean and standard deviation.

**Table S1.**
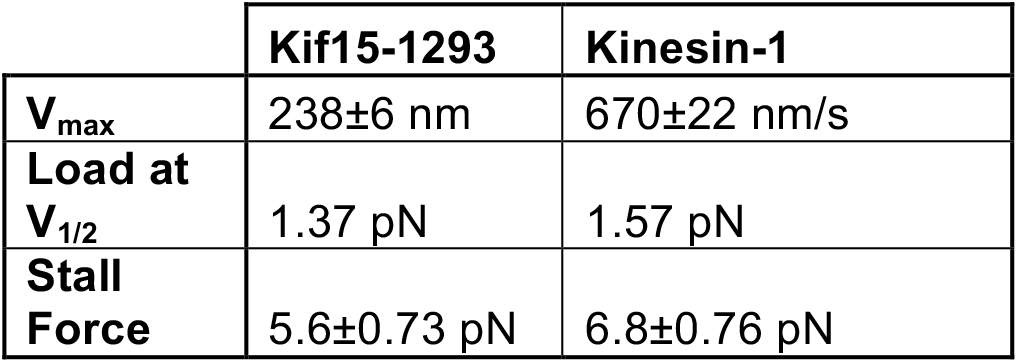

|  | **Kif15-1293** | **Kinesin-1** |
| --- | --- | --- |
| **V<sub>max</sub>** | 238±6 nm | 670±22 nm/s |
| **Load at V<sub>1/2</sub>** | 1.37 pN | 1.57 pN |
| **Stall Force** | 5.6±0.73 pN | 6.8±0.76 pN |

**Table S2.**

|  | Time in Trap Centre (s) | Tether Length (nm) |
| --- | --- | --- |
| Kinesin-1 | 3.31<br>3.04<br>3.02 | 45.5<br>38.8<br>38.3 |
| **Kinesin-1 Mean** | **3.12** | **40.8** |
| Kif15-1293 | 5.37<br>3.2<br>2.23 | 108<br>42.5<br>20.5 |
| **Kif15-1293 Mean** | **3.6** | **57** |

**Table S3.**

|  | Step Size Forward (nm) | nforward | Step Size Backward (nm) | nbackward |
| --- | --- | --- | --- | --- |
| Kif15-FL | 6.2 ± 0.1 | 314 | 6.8 ± 0.3 | 55 |
| Kif15-1293 | 7.14 ± 0.1 | 259 | 7.24 ± 0.29 | 38 |
| Kinesin-1 | 5.75 ± 0.03 | 1864 | 5.4 ± 0.1 | 583 |

### Methods

Sample preparation is as described in Drechsler et.al. (2014) and instrumentation as described in Carter and Cross (2005). Motors were non-specifically bound to polystyrene beads (550 nm, NIST calibration size standards, Polysciences, Warrington, PA, USA) by incubation in a solution of 80 mM K.Pipes, 2 mM MgSO_4_, 1 mM EGTA, 1 mM DTT, 3 mg/ml D-Glucose, 0.2 mg/ml casein and 1 μM ATP at pH 7.0. Incubation was performed for at least one hour before addition to the flow cell. Motor concentration was titrated to ensure no more than one third of beads showed any microtubule binding. A bead-binding event was defined as a bead that remained attached to the microtubule for at least 2 s after the laser trap has turned off. Each bead was tested for binding on two or more different microtubules.

Flow cells were constructed using sonicator and plasma cleaned coverslips and Dow Corning High Vacuum Grease. Taxol stabilized microtubules were diluted in assay buffer: 20 mM K.Pipes, 2 mM MgSO4, 1 mM ATP, 1 mM EGTA, 1 mM DTT, 3 mg/ml D-Glucose, 0.4 mg/ml casein, 4 μM taxol, and a glucose oxidase/catalase-based oxygen scavenging system. Diluted microtubules were introduced to the flow cell and allowed to adsorb to the surface for up to 5 minutes. The incubated bead–motor solutions were then diluted in an assay buffer and introduced to the flow cell, which was imaged immediately.

When higher ionic strengths were used for bead incubation the motors were incubated in assay buffer plus 380 mM NaCl for 30 minutes prior to incubation with beads, and the bead-motor complexes incubated in high salt buffer for at least one hour. The bead-motor solutions were then diluted in assay buffer, which remained the same throughout the experiments.

The antibody used in Figure 2 was Anti Hklp2 Antibody, Cat. Number AKIN13, from Cytoskeleton. In these experiments polystyrene beads were incubated with motors as in Carter and Cross (2005). Bead-motor complexes were then added to assay buffer containing 1mg/ml antibody and immediately introduced to the flow cell and imaged.

In order to gain an insight into the behaviour of the motor proteins at loads above stall, backwards pulls were performed. In a backwards pull the stage is moved, forcing the motor to loads above stall force. In our experiments, stage movement is automatically triggered when the motor has walked to a set load of 3pN. At this trigger point the stage is moved by piezoelectric positioning motors so that the trap centre is shifted rapidly towards the MT minus end, imposing a greater backwards load upon the motor. The load that is imposed on the motor is also pre-set, usually at either 8pN or 10pN. Whilst steps are being recorded, the stage is not moved. At these super-stall loads, motors make mostly backwards steps. Without the stage movement such high backwards loads would almost never be sampled.

Steps in bead position records were identified using a moving window t-test algorithm also described in Carter and Cross (2005). This algorithm requires the user to input a window size for the t-calculation and a t-value cut off above which a step is assigned. These were values were or Kinesin-1 data and for Kif15-FL and Kif15-1293 data.

Unbinding load experiments require the motor-bead complex to be translated past the MT at a constant rate. This was accomplished by moving the stage with piezoelectric positioning motors (Physik Instrumente (PI) 50μm x 50μm XY piezoelectric translation stage). The stage is controlled by a 12 bit DAC (Digital to

Analog Converter) allowing the stage to be stepped in 12.2 nm movements in both the x and y directions. For data collection the microtubule position and orientation were first marked and the stage then stepped at a steady rate so that the bead-motor complex was dragged along the microtubule axis. The microscope software collects data at 80-100 kHz. The primary data is sampled down to 45 kHz before storage whilst performing calibrations and to 22 kHz whilst collecting motor stepping data.

The laser trap calibration was performed daily, stiffness was assigned as the average value of stiffness determined using three different methods; Equipartition, Stokes and drag force.

The rate at which load is applied can make a large difference to the distribution of measured unbinding loads. When comparing data sets the data were cropped so that for each set of data the loading rates were similar (5±0.5 pN/s) and only a small range of loading rates was used (Figure S4D). In cases where the bead did not return to the trap centre after a detachment (a backward slip greater than 16nm) the second detachment is neglected.

## Bibliography

Andreasson, J. O., B. Milic, G. Y. Chen, N. R. Guydosh, W. O. Hancock and S. M. Block (2015). “Examining kinesin processivity within a general gating framework.” Elife 4.

Balchand, S. K., B. J. Mann, J. Titus, J. L. Ross and P. Wadsworth (2015). “TPX2 Inhibits Eg5 by Interactions with Both Motor and Microtubule.” J Biol Chem 290(28): 17367–17379.

Carter, N. J. and R. A. Cross (2005). “Mechanics of the kinesin step.” Nature 435(7040): 308–312.

Chen, G. Y., K. J. Mickolajczyk and W. O. Hancock (2016). “The Kinesin-5 Chemomechanical Cycle Is Dominated by a Two-heads-bound State.” J Biol Chem 291(39): 20283–20294.

Clancy, B. E., W. M. Behnke-Parks, J. O. Andreasson, S. S. Rosenfeld and S. M. Block (2011). “A universal pathway for kinesin stepping.” Nat Struct Mol Biol 18(9): 1020–1027.

Cross, R. A. and A. McAinsh (2014). “Prime movers: the mechanochemistry of mitotic kinesins.” Nat Rev Mol Cell Biol 15(4): 257–271.

Drechsler, H. and A. D. McAinsh (2016). “Kinesin-12 motors cooperate to suppress microtubule catastrophes and drive the formation of parallel microtubule bundles.” Proc Natl Acad Sci U S A 113(12): E1635–1644.

Drechsler, H., T. McHugh, M. R. Singleton, N. J. Carter and A. D. McAinsh (2014). “The Kinesin-12 Kif15 is a processive track-switching tetramer.” Elife 3: e01724.

Hancock, W. O. and J. Howard (1999). “Kinesin’s processivity results from mechanical and chemical coordination between the ATP hydrolysis cycles of the two motor domains.” Proc Natl Acad Sci U S A 96(23): 13147–13152.

Korneev, M. J., S. Lakamper and C. F. Schmidt (2007). “Load-dependent release limits the processive stepping of the tetrameric Eg5 motor.” Eur Biophys J 36(6): 675–681.

Mann, B. J., S. K. Balchand and P. Wadsworth (2017). “Regulation of Kif15 localization and motility by the C-terminus of TPX2 and microtubule dynamics.” Mol Biol Cell 28(1): 65–75.

Rath, O. and F. Kozielski (2012). “Kinesins and cancer.” Nat Rev Cancer 12(8): 527–539.

Saunders, A. M., J. Powers, S. Strome and W. M. Saxton (2007). “Kinesin-5 acts as a brake in anaphase spindle elongation.” Curr Biol 17(12): R453–454.

Scholey, J. E., S. Nithianantham, J. M. Scholey and J. Al-Bassam (2014). “Structural basis for the assembly of the mitotic motor Kinesin-5 into bipolar tetramers.” Elife 3: e02217.

Shimamoto, Y., S. Forth and T. M. Kapoor (2015). “Measuring Pushing and Braking Forces Generated by Ensembles of Kinesin-5 Crosslinking Two Microtubules.” Dev Cell 34(6): 669–681.

Sturgill, E. G., D. K. Das, Y. Takizawa, Y. Shin, S. E. Collier, M. D. Ohi, W. Hwang, M. J. Lang and R. Ohi (2014). “Kinesin-12 Kif15 targets kinetochore fibers through an intrinsic two-step mechanism.” Curr Biol 24(19): 2307–2313.

Sturgill, E. G. and R. Ohi (2013). “Kinesin-12 differentially affects spindle assembly depending on its microtubule substrate.” Curr Biol 23(14): 1280–1290.

Tanenbaum, M. E., L. Macurek, A. Janssen, E. F. Geers, M. Alvarez-Fernandez and R. H. Medema (2009). “Kif15 cooperates with eg5 to promote bipolar spindle assembly.” Curr Biol 19(20): 1703–1711.

Uemura, S., K. Kawaguchi, J. Yajima, M. Edamatsu, Y. Y. Toyoshima and S. Ishiwata (2002). “Kinesin-microtubule binding depends on both nucleotide state and loading direction.” Proc Natl Acad Sci U S A 99(9): 5977–5981.

Valentine, M. T. and S. M. Block (2009). “Force and premature binding of ADP can regulate the processivity of individual Eg5 dimers.” Biophys J 97(6): 1671–1677.

Valentine, M. T., P. M. Fordyce, T. C. Krzysiak, S. P. Gilbert and S. M. Block (2006). “Individual dimers of the mitotic kinesin motor Eg5 step processively and support substantial loads in vitro.” Nat Cell Biol 8(5): 470–476.

van Heesbeen, R. G., M. E. Tanenbaum and R. H. Medema (2014). “Balanced activity of three mitotic motors is required for bipolar spindle assembly and chromosome segregation.” Cell Rep 8(4): 948–956.

Vanneste, D., M. Takagi, N. Imamoto and I. Vernos (2009). “The role of Hklp2 in the stabilization and maintenance of spindle bipolarity.” Curr Biol 19(20): 1712–1717.

